# Extending the breeder’s equation to take aim at the Target Population of Environments

**DOI:** 10.1101/2022.12.13.520360

**Authors:** Mark Cooper, Owen Powell, Carla Gho, Tom Tang, Carlos Messina

## Abstract

A major focus for genomic prediction has been on improving trait prediction accuracy using combinations of algorithms and the training data sets available from plant breeding multi-environment trials (METs). Any improvements in prediction accuracy are viewed as pathways to improve traits in the reference population of genotypes and product performance in the target population of environments (TPE). To realise these breeding outcomes there must be a positive MET-TPE relationship that provides consistency between the trait variation expressed within the MET data sets that are used to train the genome-to-phenome (*G2P*) model for applications of genomic prediction and the realised trait and performance differences in the TPE for the genotypes that are the prediction targets. The strength of this MET-TPE relationship is usually assumed to be high, however it is rarely quantified. To date investigations of genomic prediction methods have not given adequate attention to quantifying the structure of the TPE and the MET-TPE relationship and its potential impact on training the *G2P* model for applications of genomic prediction to accelerate breeding outcomes for the on-farm TPE. We provide a perspective on the importance of the MET-TPE relationship as a key component for the design of genomic prediction methods to realize improved rates of genetic gain for the target yield, quality, stress tolerance and yield stability traits in the on-farm TPE.

## 2) Introduction

Plant breeding is grounded in prediction (Goldman, 2000; Duvick, 2001; Cooper et al., 2014a; Voss-Fels et al., 2019). Plant breeding programs are the operational implementation of coordinated sequences of prediction methods, organised to continuously create, evaluate and select new genotypes over multiple breeding program cycles (Duvick et al., 2004; Cobb et al., 2019; Technow et al., 2021). The cycles are designed to iteratively improve on the outcomes from previous cycles. Breeding objectives are framed to develop product outcomes (varieties, hybrids, clones, populations). These products are to be used by farmers within the Genotype-by-Environment-by-Management (GxExM) context of agricultural systems of the target population of environments (TPE); which includes the biophysical environment and the agronomic management practices adopted by farmers (Cooper et al., 2020, 2021; Kholová et al., 2021; Ronanki et al., 2022). Through successful adoption and use of the improved products by farmers, breeding programs can improve food productivity and so contribute to enhanced global food security. However, there are many persistent gaps documented between the current levels of crop productivity in agricultural systems and the targets required to achieve food security. Thus, there is continued interest in improving the design of breeding programs to target the creation of new products to help close yield gaps (Cooper et al., 2020; Messina et al., 2022a). Application of genomic prediction technologies has emerged as a major theme of breeding program design in the 21^st^ Century (Meuwissen et al., 2001; Heffner et al., 2009; Cooper et al., 2014a; Voss-Fels et al., 2019; Rogers et al., 2021; Varshney et al., 2021). Here we discuss and extend the “breeder’s equation” as a framework to help evaluate opportunities to enhance genomic breeding outcomes through enhanced design of METs to provide the relevant training data sets with the required MET-TPE alignment (Cooper et al., 2014a, b; Gaffney et al., 2015; González-Barrios et al., 2019; Rogers et al., 2021; Smith et al., 2021a, b). Attention to improve the MET-TPE alignment as a criterion in the design of MET training data sets supports effective use of environmental covariates, crop models and high-throughput phenotyping in combination with genome-to-phenome (*G2P*) modelling algorithms to enhance genomic prediction for the TPE (Cooper et al., 2014a, b; Gaffney et al., 2015; Messina et al., 2018; Diepenbrock et al., 2021).

The basic form of the “breeder’s equation” provides a framework to predict the response to selection (Δ*G*) from one cycle (*L*) of a breeding program, following application of a selection strategy. Here we consider selection strategies that incorporate applications of genomic prediction (Heffner et al., 2009; Cooper et al., 2014a; Voss-Fels et al., 2019). Selection pressure is implemented by applying truncation selection to the distributions of observed or predicted values for one or more traits within the reference population of genotypes (RPG) of a breeding program; for example, selection to increase crop yield, improve grain quality and improve abiotic and biotic stress tolerances to reduce the extent of yield losses due to the occurrence of the frequent stresses in the TPE (Chenu et al., 2011; Kholová et al., 2013; Hajjarpoor et al., 2021; Messina et al., 2022a). The structure of the breeder’s equation has a long history in animal and plant breeding (Lush, 1937; Hallauer and Miranda, 1988; Nyquist and Baker, 1991; Comstock, 1996) and is frequently used as a quantitative framework for the design and optimisation of crop breeding programs (Araus and Cairns, 2014; Cobb et al., 2019; Kholová et al., 2021; Cooper and Messina, 2022). For applications of genomic prediction, a common form of the breeder’s equation is given as:

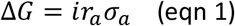

Where *i* represents the selection differential applied to the selection units, based on the trait variation within the RPG, *r*_*a*_ represents the prediction accuracy for breeding values for the selection units within the RPG, and *σ*_*a*_ represents the additive genetic variation among the selection units within the RPG for the traits that are targeted for improvement by selection. For genomic breeding, the quantification of prediction accuracy *r*_*a*_ is based on *G2P* models for traits that are constructed using suitable training data sets. These training data sets are created algorithmically using the genetic fingerprints and trait phenotypes for the genotypes included in breeding multi-environment trials (METs) used as training data sets. The foundation of the MET training data sets is typically based on data collected from the relevant stages of the breeding program (Cooper et al., 2014a; Smith et al., 2021a). Characterisations of the sample of environments present in the MET can be used to create environmental predictors to be included in the *G2P* model to adjust genomic predictions to account for effects of GxE interactions (Jarquín et al., 2014; Messina et al., 2018; de los Campos et al., 2020; Diepenbrock et al., 2021). Importantly, the samples of environments included in the METs are considered to represent the environmental composition of the TPE (Comstock and Moll, 1963; Nyquist and Baker 1991; Cooper and DeLacy, 1994). The environmental composition of the METs can be augmented in many ways using specifically designed field-based and controlled-environment experiments (Cooper et al., 1995, 1997, 2014a, b; Campos et al., 2004; van Eeuwijk et al., 2019; Langstroff et al., 2022; Cooper and Messina, 2022). Many assumptions are made when applying the breeder’s equation, as represented by equation (1). We consider some of these assumptions in more detail as they relate to the prediction of response to selection for improved on-farm performance within the TPE. In particular we focus on the influence of the MET-TPE relationship in the presence of GxE interactions within the TPE of the breeding program and use this as the basis for deriving the extended breeder’s equation introduced here.

### 3.1 Extending the breeder’s equation to take aim at the TPE

The breeder’s equation, as represented in equation (1), quantifies the per cycle rate of change of the trait mean value for the RPG (Nyquist and Baker, 1991; Cobb et al. 2019). However, the breeder’s equation does not explicitly quantify the directionality of the change in trait values relative to their requirements for the TPE. To enable efficient design of a breeding program, targeted on creation of new products to close on-farm yield gaps, it is desirable to have a form of the breeder’s equation that includes both the rate and the directionality components of genetic gain for the TPE. Applying correlated response selection theory (Falconer, 1952; Cooper and DeLacy, 1994; Rogers et al., 2021; Cooper and Messina, 2022), we provide an extended form of the breeder’s equation that combines both the rate and directionality components of trait change under the influence of selection. Considering the environmental composition of the MET to be a sample of the environmental composition of the TPE (*MET* ∈ *TPE*), an equation for trait genetic gain within the TPE, based on selection decisions made using predictions from *G2P* trait information obtained from METs (Δ*G* (_*MET,TPE*)_), can be given as:

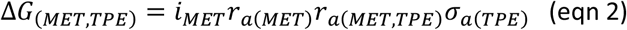

Two of the terms in equation (2) are equivalent to terms in equation (1): *i*_*MET*_ is the selection differential applied to phenotypic and *G2P* prediction information obtained from analyses of the MET training data sets, as for *i* in equation (1), *r*_*a*(*MET*)_ is the prediction accuracy for the selection units based on applications of the training data available from the MET, as for *r*_*a*_ in equation (1). In equation (2) the *σ*_*a*_ term of equation (1) is replaced by the product of two terms *r*_*a*(*MET,TPE*)_ and *σ*_*a*(*TPE*)_. The term *r*_*a*(*MET,TPE*)_ is the genetic correlation between the additive genetic effects estimated by applying *G2P* models developed using the MET training data sets, and the additive genetic effects for the trait targets required for realised trait performance in the TPE. The term *σ*_*a*(*TPE*)_ represents the additive genetic variation for the traits within the TPE. Additional forms of equation (2) can be given, for example for prediction at the level of the total genotypic trait performance level. Equally equation (2) can be further extended to examine the contributions of quantitative trait loci (QTL) and combinations of haplotypes and specific QTL to the additive or total genotypic variance for multiple traits in the RPG for the TPE.

Applying the extended form of the breeder’s equation given in equation (2), statements can be made regarding the design of genomic prediction strategies based on applications of equation (1).

- Firstly, if the environmental composition of the MET is an accurate sample of the environmental composition of the TPE then it can be expected that *r*_*a*(*MET,TPE*)_ → 1 and equations (1) and (2) will converge to the same form of the breeder’s equation, as given in equation (1); in this case the *σ*_*a*_ of equation (1) converges to the *σ*_*a*(*TPE*)_ of equation (2). However, if there is GxE interaction and divergence in environmental composition between the MET and the TPE, *r*_*a*(*MET,TPE*)_ < 1 can occur, diminishing prediction accuracy for the TPE. Under such circumstances it can be expected that realised genetic gain in the TPE will be lower than predicted when based on studies confined to pursuing *G2P* modelling algorithms for improved prediction accuracy within the bounds of the MET training data sets; in this case the *σ*_*a*_ of equation (1) can diverge from the *σ*_*a*(*TPE*)_ of equation (2). Whenever there is historical evidence that realised genetic gains in the on-farm TPE are lower than the predicted gains, the magnitude of *r*_*a*(*MET,TPE*)_ should be investigated to quantify its potential impact on the expected realised prediction accuracy that can be achieved in the TPE based on prediction accuracy derived from the training data available through the MET.
- Secondly, whenever there is evidence of GxE interactions within the TPE, including GxExM interactions, and there is the potential for divergence between the environmental composition and trait data obtained from current METs and those expected for the future TPE, as is often projected for the influences of climate change, the extended form of the breeder’s equation (2) provides a more appropriate framework than equation (1) for quantifying the impact of such changes on the design and optimisation of prediction-based breeding strategies.
- Thirdly, for long-term breeding programs, consideration should be given to characterisation of the TPE and the design of MET experiments to obtain empirical estimates of the genetic correlation *r*_*a*(*MET,TPE*)_ and determination of the genetic and environmental factors contributing to *r*_*a*(*MET,TPE*)_ < 1. The effects of climate change on the environmental composition of the TPE and associated changes in trait contributions to yield and GxE interactions for current and future cropping systems represents one clear area for urgent consideration in the design of METs to address the MET-TPE alignment (Braun et al., 2010; Chapman et al., 2012; Lobell et al., 2015; Ceccarelli and Grando, 2020; IPCC, 2021; Snowdon et al., 2021; Bustos-Korts et al., 2021; Cooper et al., 2021, Cooper and Messina, 2022).

To demonstrate the implications of GxE interactions on realized genetic gain in the on-farm TPE we consider two examples of the application of the extended form of the breeder’s equation to investigate the MET-TPE alignment and its potential impact on the *r*_*a*(*MET,TPE*)_ component of equation (2). The first considers a familiar theoretical example from the study of crossover GxE interactions (Haldane, 1947; Ceccarelli, 1989, 1994; Cooper and DeLacy, 1994; van Eeuwijk et al., 2001). The second considers an empirical example based on a previously published MET-TPE data set for wheat in Australia (Cooper et al., 1995, 1997, 2001). The wheat example was previously used to investigate the implications of GxE interactions for grain yield in the TPE, and also the MET-TPE relationship for design of METs to accelerate genetic gain for yield from wheat breeding in a TPE where complex GxE interactions for grain yield are ubiquitous (Brennan et al., 1981; Cooper and DeLacy, 1994; Cooper et al., 1995, 1997, 2001; Basford and Cooper, 1998, Chenu et al., 2011; Lobell et al., 2015; Bustos-Korts et al., 2021; Resende et al., 2021).

### 3.2 Investigating the MET-TPE Alignment: Theoretical Example

Theoretical and empirical considerations of the influences of GxE interactions for breeding have consistently emphasised the importance of crossover GxE interactions (Figure 1a; Haldane 1947; Ceccarelli 1989, 1994; Cooper and DeLacy 1994; Cooper et al. 2021; Rogers et al. 2021; Smith et al. 2021a, b). Examples of such crossover interactions in breeding METs have been demonstrated at the genotypic (Cooper et al. 1995, 1997, van Eeuwijk et al. 2001, Xiong et al. 2021; Smith et al. 2021b) and QTL levels (Boer et al. 2007, Millet et al. 2019). For the theoretical example of crossover GxE interactions shown in Figure 1a, the yield performance responses for two genotypes (G2 and G8) in two environments (Env_1 and Env_2) are considered. The potential impact of the crossover interactions depicted in Figure 1a on selection decisions can be examined using equation (2) by considering the influence of changes in the frequency of occurrence of the two environments within the both the MET and TPE on the genetic correlation *r*_*a*(*MET,TPE*)_ term from equation (2). Here we consider the genotypic correlation *r*_*g*(*MET,TPE*)_ between weighted average yield of the two genotypes between the MET and the TPE, where the weights are based on the frequencies of occurrence of the two environments in the MET and the TPE (Podlich et al. 1999). This provides a simulated scan of the range of possible MET-TPE alignment scenarios based on the potential range in frequency of occurrence of the two environments within the MET and the TPE.

**Figure 1.**
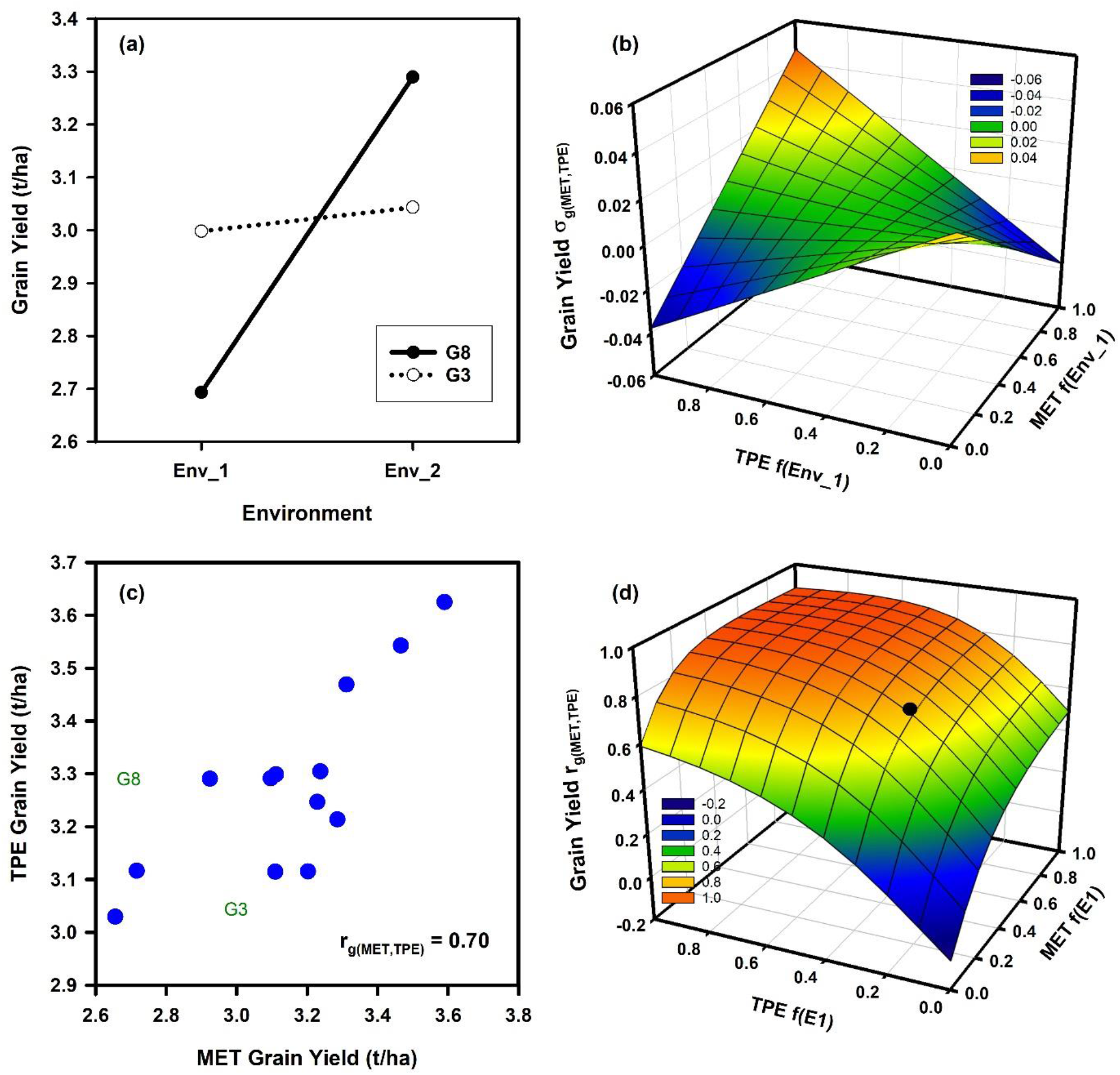
Two examples of the potential influences of Genotype by Environment (GxE) interactions for grain yield on the expected genetic correlation between the average genotype performance in a multi-environment trial (MET) and the target population of environments (TPE) as the frequencies of environment types change between the sample of environments obtained in the MET and their presence in the TPE: (a) Schematic yield reaction-norms for two wheat genotypes (G3, G8) in two environments (Env_1, Env_2) demonstrating crossover GxE interaction; (b) Response surface of the expected genotypic covariance *σ*_*g*(*MET,TPE*)_ between average genotype yield performance in a MET and in the TPE as the frequencies of the two environments (Env_1, Env_2) change within the MET and TPE; (c) Scatter plot of the average grain yield for 15 wheat genotypes based on two independent sets of environments representing both the MET and the TPE; (d) Response surface of the expected genotypic correlation, *r*_*g*(*MET,TPE*)_ from equation (2), between average genotype yield performance in a MET and in the TPE as the frequencies of two environment-types (E1 = Mild water deficit, E2 = Severe water-deficit) change within the MET and TPE data sets. The filled symbol on the response surface positions the estimate of *r*_*g*(*MET,TPE*)_ for the grain yield data shown in sub-figure 2c (MET f(E1) = 0.41, TPE f(E1) = 0.31, *r*_*g*(*MET,TPE*)_ = 0.70). Data for grain yield estimates were obtained from the study reported by Cooper et al. (1997).

In Figure 1b the genotypic covariance *σ*_*g*(*MET,TPE*)_ of the average performance of the two genotypes in the MET and the TPE is plotted against the frequency of Env_1 in the MET and the TPE. The genotypic covariance is the numerator of the genetic correlation *r*_*g*(*MET,TPE*)_ term of equation (2) and is used here in place of *r*_*g*(*MET,TPE*)_ to smooth out the response surface for illustration purposes. The shape of the response surface for the genotypic covariance (Figure 1b) fluctuates between negative and positive values depending on the frequency of occurrence of both environments in the MET and the TPE. Two aspects are noted.

- Firstly, when the frequencies of both environments are close to 0.5 in the MET or TPE the genetic covariance, and thus the genetic correlation *r*_*g*(*MET,TPE*)_, approaches 0 (Figure 1b). In such situations selection decisions will require direct investigation of the GxE interactions and consideration of how to target breeding for both environments instead of selection for average performance in the MET to improve average performance in the TPE, as simulated here (Figure 1a).
- Secondly, as the frequencies of the environments within the MET and the TPE deviate from 0.5 towards 1.0 for Env_1 and towards 0.0 for Env_2, or towards 0.0 for Env_1 and towards 1.0 for Env_2, then the influence of the MET-TPE alignment becomes increasingly important. When there is MET-TPE alignment of the environment frequencies the genotypic covariance is positive and the crossover GxE interaction is less problematic for selection decisions (Figure 1b). However, if there is poor MET-TPE alignment of the environment frequencies, for example a high frequency of Env_1 in the MET when Env_1 actually has a low frequency in the TPE, then the genotypic covariance can become negative (Figure 1b). In this situation selection based on the information obtained from the MET will result in poor selection decisions that are not aligned with the needs of the TPE, even if a high prediction accuracy, based on the value of *r*_*a*_ from equation (1) and of *r*_*a*(*MET*)_ from equation (2), is demonstrated for any prediction method within the confines of the MET training data set.

### 3.3 Investigating the MET-TPE Alignment: Empirical Example

Building on the theoretical example (Figure 1a, b), we apply the extended breeder’s equation to quantify the impact of the MET-TPE alignment for an empirical example by estimating the genotypic correlation *r*_*g*(*MET,TPE*)_ term of equation (2) for a range of wheat MET-TPE alignment scenarios for north-eastern Australia (Figure 1c, d). We utilise grain yield data available from a previously published wheat data set (Cooper et al., 1995, 1997, 2001). The example provides grain yield data for 15 genotypes and 53 environments. Importantly, for current considerations, the 53 environments were previously organised to represent a breeding MET (27 environments) and the TPE (26 environments) for the north-eastern region of the Australian wheat belt (Brennan et al., 1981; Cooper et al., 1995, 1997; Chenu et al., 2011). The MET was specifically designed in an attempt to represent the current understanding of GxE interactions and MET-TPE alignment scenarios for the wheat breeding program at that time. The set of 15 genotypes was chosen to represent groupings of key germplasm from the reference population of genotypes for the wheat breeding program (Cooper and DeLacy, 1994; Cooper et al., 1995, 1997, 2001). Further, we identify that the data for the two genotypes (G2 and G8), used to illustrate crossover GxE interactions in the theoretical example (Figure 1a), were chosen from the larger set of 15 genotypes included in the empirical example (Figure 1c). Also, the two environments (Env_1 and Env_2) used in the theoretical example were taken from the empirical example. Thus, the numerical values for the example of crossover GxE interaction for grain yield (Figure 1a) used for the theoretical investigations (Figure 1b) were representative of important crossover GxE interactions under consideration within the target breeding program, as considered in the empirical example (Figure 1c, d; Brennan et al., 1981; Cooper and DeLacy, 1994; Basford and Cooper, 1998; Cooper et al., 2001).

Improving grain yield stability for the TPE of the north-eastern region of the Australian wheat-belt was a primary objective of the wheat breeding program at that time. A weighted selection strategy, combined with field-based managed-environments, was developed to account for GxE interactions (Cooper et al., 1995, 1997, 2001; Podlich et al., 1999). Spatial and temporal variability for water availability was identified as primary driver of grain yield variation within the TPE, and drought was considered to be a major source of crossover GxE interactions for grain yield. Thus, the environments included in the MET were managed to sample a gradient of water availability scenarios, ranging from severe drought to water-sufficient environments, by managing combinations of irrigation and nitrogen inputs at a restricted number of locations. The TPE set of environments was designed by sampling a range of water availability scenarios from a wider range of locations and years within the north-eastern region of Australia. The objective was to design a MET for the stages of the wheat breeding program that could be consistently managed at a few locations to provide a stratified sample of the range of water availability environments expected within the TPE (Cooper et al., 1995, 1997, 2001).

Grain yield GxE interactions were previously identified for both the MET and TPE data sets (Cooper et al., 1995, 1997, 2001). Crossover GxE interactions were frequent (Figure 1a; Cooper and DeLacy, 1994). For the purposes of demonstrating an application of equation (2) to the empirical wheat example, the prior envirotyping was used to identify two groups of environment-types for both the MET and TPE sets; environment-type 1 (E1) characterised by mild water-deficits, and environment-type 2 (E2) characterised by severe water-deficits. There were GxE interactions between the two environment-types within the MET and TPE sets (Figure 2; Cooper et al., 1995, 1997, 2001). There was a moderate to weak positive genotypic correlation for grain yield variation among the 15 genotypes between both environment-types E1 and E2 for the MET (Figure 2a) and TPE (Figure 2b). Importantly, for interpretation of the genotypic correlation *r*_*g*(*MET,TPE*)_ between the MET and TPE (Figure 1d), the genotypic correlation for grain yield variation between the mild stress environment-type E1 was positive and strong between the MET and the TPE (Figure 2c). However, there was no relationship for grain yield variation between the severe drought stress environment-type E2 between the MET and the TPE (Figure 2d). The details of the lack of relationship for environment-type E2 are discussed in detail elsewhere (Cooper et al., 1995, 1997). In summary the MET was designed to focus on the expected water availability gradient in the absence of other abiotic and biotic stresses that could also occur within the TPE. Occurrences of these other abiotic and biotic stresses within the TPE set were interpreted to be contributing factors to the low relationship observed for severe drought stress environment-type E2 between the MET and TPE (Figure 2d). In the absence of the drought stress for environment-type E1 these other abiotic and biotic stresses were less influential on the genotypic correlation for grain yield (Figure 2c).

**Figure 2.**
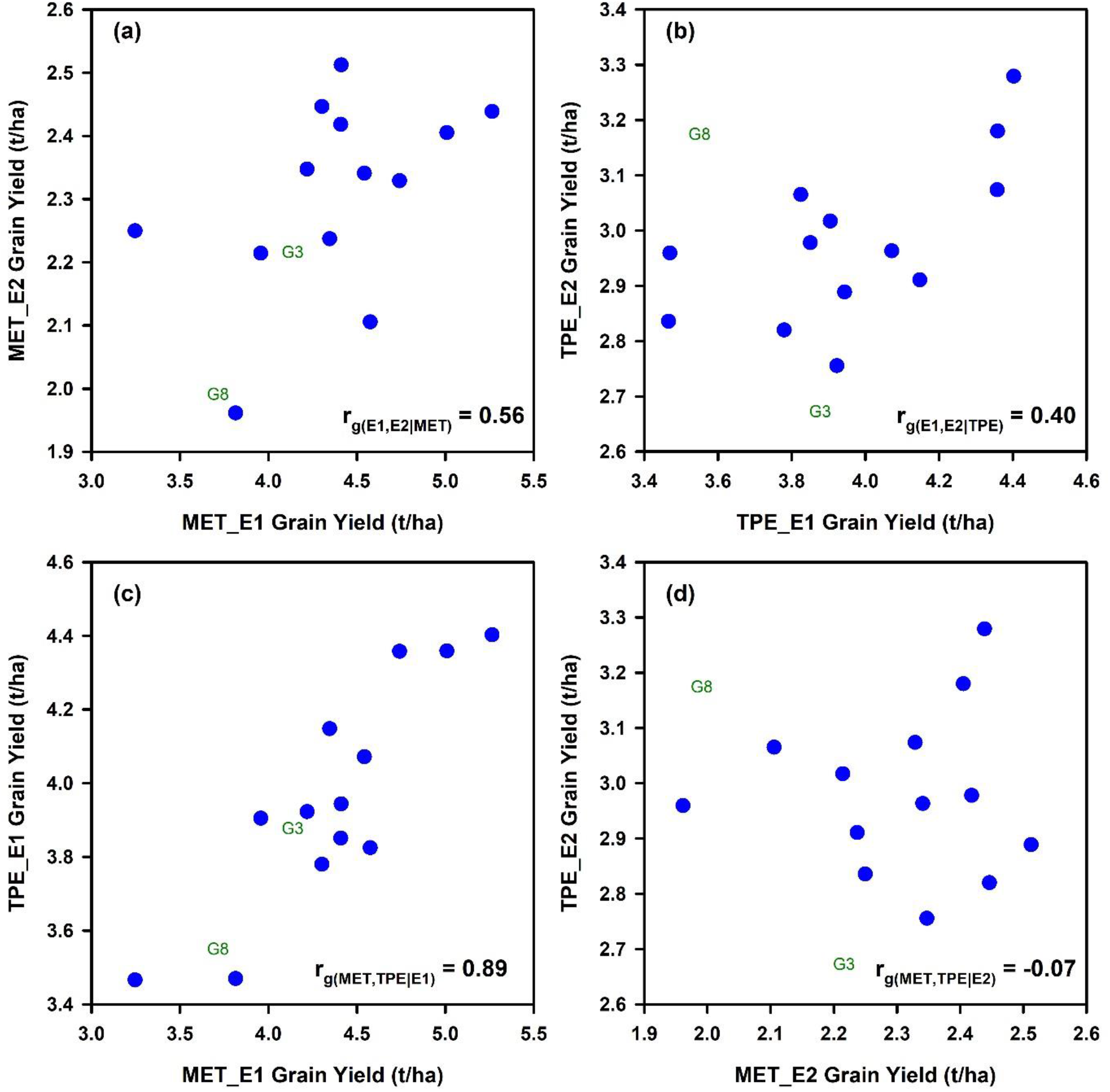
Scatter diagrams comparing average grain yield predicted for 15 wheat genotypes for two environment-types (E1 = Mild water deficit, E2 = Severe water-deficit) obtained from independent data sets representing a multienvironment trial (MET) and the target population of environments (TPE): (a) Comparison between grain yield predicted for environment-types E1 and E2 in the MET data set, *r*_*g*(*E*1,*E*2|*MET*)_; (b) Comparison between grain yield predicted for environment-types E1 and E2 in the TPE data set, *r*_*g*(*E*1,*E*2|*TPE*)_; (c) Comparison of grain yield predicted for environment-type E1 between the MET and the TPE data sets, *r*_*g*(*MET,TPE*|*E*1)_; (d) Comparison of grain yield predicted for environment-type E2 between the MET and the TPE data sets, *r*_*g*(*MET,TPE*|*E*2)_. Data for grain yield predictions were obtained from the study reported by Cooper et al. (1997).

For purposes of demonstrating an application of the extended breeder’s equation to the wheat MET-TPE data set (Figure 1c, d) it is sufficient to note that there was GxE interaction for grain yield between Environment-types E1 and E2 in both the MET (Figure 2a) and the TPE (Figure 2b) data sets and that there was positive predictability between the MET and TPE sets for environment-type E1 (Figure 2c), but no predictability for environment-type E2 (Figure 2d). Using this level of envirotyping we can simulate the influence of changes in the MET-TPE alignment on *r*_*g*(*MET,TPE*)_ and prediction of average grain yield in the TPE based on average grain yield estimated from the MET (Figure 1d). Following the same procedures applied to the theoretical example (Figure 1a, b), the potential range of MET-TPE alignment scenarios was simulated by changing the frequencies of environment-types E1 and E2 within the MET and the TPE in steps of 0.1 from 0.0 to 1.0, calculating the weighted average grain yield of the 15 genotypes for both the MET and TPE, taking into consideration the frequencies of both environment-types, and calculating the genotypic correlation *r*_*g*(*MET,TPE*)_ between the estimates of weighted average grain yield for the 15 genotypes between the MET and TPE for all MET-TPE alignment combinations. We then plotted the *r*_*g*(*MET,TPE*)_ against the frequency of environment-type E1 in the MET and TPE to generate a simulated *r*_*g*(*MET,TPE*)_ genotypic correlation response surface for all MET-TPE alignment configurations (Figure 1d).

The genotypic correlation *r*_*g*(*MET,TPE*)_ between the simulated MET and TPE alignments ranged from a high value of 0.90 to a low value of -0.07 (Figure 1d). The *r*_*g*(*MET,TPE*)_ response surface for the wheat example has interesting features. Firstly, there is a relatively broad plateau of high *r*_*g*(*MET,TPE*)_ values for many of the MET-TPE alignment scenarios. This plateau of high *r*_*g*(*MET,TPE*)_ values occurred for scenarios where the frequency of the water-sufficient environment-type E1 was higher than 0.5 in both the MET and TPE (Figure 1d), taking advantage of the high predictability between environment-type E1 in the MET and TPE (Figure 2c). Secondly, when the frequency of environment-type E1 falls below 0.5 in the MET or TPE, and therefore the frequency of the water-limited environment-types E2 increases above 0.5, the *r*_*g*(*MET,TPE*)_ is degraded from the high levels of the plateau (Figure 1d), reflecting the increased influence of the poor predictability between the MET and TPE for the water-limited environment-type E2 (Figure 2d). This impact of the MET-TPE alignment on predictability for performance in the TPE using MET results will apply to all levels of prediction, including genomic prediction, phenotypic prediction and combined prediction approaches.

For the specific environment-type configuration realised for the empirical example (Figure 2), the estimate of *r*_*g*(*MET,TPE*)_ for prediction of average grain yield for the TPE based on average gain yield obtained for the MET was intermediate (Figure 1c); *r*_*g*(*MET,TPE*)_ = 0.70 for MET *f(E1)* = 0.41, *f(E2)* = 0.59 and for TPE *f(E1)* = 0.31, *f(E2)* = 0.69. Thus, the MET-TPE alignment for the empirical example was located on the *r*_*g*(*MET,TPE*)_ response surface (Figure 1d) slightly off of the plateau of higher *r*_*g*(*MET,TPE*)_ levels, but still above the precipice where the *r*_*g*(*MET,TPE*)_ value is severely degraded. This empirical realisation of MET-TPE alignment is just one of the many possible scenarios that can occur as the frequencies of environment-types change between the MET and the TPE (Figure 1d).

The empirical wheat example (Figures 1, 2) was used to demonstrate the utility of the extended form of the breeder’s equation for applications in prediction-based breeding. Here we have emphasised the use of the extended breeder’s equation as a useful framework to guide the design MET data sets for training *G2P* models for applications of genomic prediction and genomic selection at different stages of a breeding program to take aim at the TPE (Cooper et al., 2014a, b; Gaffney et al. 2015; Messina et al. 2022a). Many other possible prediction scenarios can also be investigated, and these will be the subject of future research.

## 4) Discussion

Design of breeding programs, and crop improvement strategies in general, to take aim at the crop productivity requirements of the TPE is critical to both accelerate and achieve realised genetic gain on-farm that contributes to closing yield gaps (Messina et al., 2022a), improving global food security (Cooper et al., 2021; Kholová et al., 2021; Rogers et al., 2021), and the many other requirements for sustainable agricultural systems (Messina et al., 2022b). However, in most considerations of breeding program design and optimisation there is no direct connection between the optimisation considerations that use the framework of the breeder’s equation, as in equation (1), and the understanding of the TPE. Thus, there is often a disconnect between the attention to rate of genetic gain, the directionality of the breeding program and its MET-TPE alignment with the requirements of the on-farm TPE. In the presence of GxE interactions and low *r*_*a*(*MET,TPE*)_ this MET-TPE disconnect can result in low realised genetic gain under the on-farm conditions of the TPE, even when high prediction accuracy, based on *r*_*a*_ in equation (1) or more explicitly *r*_*a*(*MET*)_ in equation (2), is demonstrated for genomic prediction methods evaluated within the confines of the MET. The extended form of the breeder’s equation, introduced here as equation (2), provides a framework to remove this disconnect and to support design of prediction-based breeding strategies that take aim at the TPE by emphasising the influence of the MET-TPE alignment on realised genetic gain for the on-farm TPE (Cooper et al., 2014a, b; Gaffney et al. 2015; Messina et al. 2022a). Here we demonstrated such application of the extended breeder’s equation framework through investigation of *r*_*a*(*MET,TPE*)_, rather than assuming *r*_*a*(*MET,TPE*)_ = 1, as is the case for the traditional form of the breeder’s equation.

We have introduced and demonstrated the utility of the extended form of the breeder’s equation through applications to a theoretical and empirical example. In summary the following key points were presented.

### Theoretical considerations

We extended the breeder’s equation, introducing the genetic correlation *r*_*a*(*MET,TPE*)_ to explicitly incorporate and quantify the relationship between a MET and the TPE, as a framework for designing METs to take aim at the TPE. Three further considerations are important: (1) the traditional form of the breeder’s equation assumes that the genetic correlation *r*_*a*(*MET,TPE*)_ = 1; (2) in the presence of GxE interactions the genetic correlation *r*_*a*(*MET,TPE*)_ can be decomposed to take into account the genetic variance-covariance structure among the environment-types within the TPE (Cooper and DeLacy, 1994; van Eeuwijk et al., 2001; Smith et al., 2005, 2021a, b; Rogers et al., 2021); and (3) the genetic correlation *r*_*a*(*MET,TPE*)_ can be applied to the continuum of selection units of interest to breeders, extending from the level of sequence information, accounting for QTL and chromosomal haplotypes, to total multi-trait, multi-QTL predicted genotypic performance or breeding value obtained for any *G2P* model that is derived from relevant training data sets that can be generated from METs together with augmented data sources from specialised phenotyping facilities (Cooper et al., 2014a,b; Gaffney et al., 2015; Diepenbrock et al., 2021).

Taking aim at specific target environment-types, for example specific biotic or abiotic stresses, is not uncommon in plant breeding (Blum, 1988; Millet et al., 2019). However, taking aim at the TPE as a mixture of environment-types (Podlich et al., 1999; Cooper et al., 2014a,b; Gaffney et al., 2015; Rogers et al., 2021; Smith et al., 2021a, b; Messina et al., 2022a,b) is much less common than taking aim at specific environment-types. Taking aim at the TPE requires detailed consideration of the mixture of target environment-types within the TPE (Chapman et al., 2000; Chenu et al., 2011; Kholová et al., 2013; Cooper et al., 2014a, b; Lobell et al., 2015; Hajjarpoor et al., 2021; Resende et al., 2021), the extent of GxE interactions between environment-types (Figure 2) and the details of the genetic variance-covariance structure among the environment-types, and appropriate attention to weighting the sources of *G2P* information for traits, that is available from the environment-types sampled in the MET training data sets, by their frequencies of occurrence and relative importance in the TPE (Podlich et al., 1999; Cooper et al., 2014a, b; Gaffney et al., 2015; Messina et al., 2018; Smith et al., 2021b).

### Empirical considerations

We demonstrated the application of the extended form of the breeder’s equation by applying it to a grain yield data set designed for a wheat breeding program, where the environments had previously been grouped into MET and TPE sets with a characterisation of the different environment-types in both the MET and TPE sets (Figures 1 and 2; Cooper et al., 1995, 1997, 2001). This prior characterisation of environment-types and the MET-TPE alignment was conducted prior to the more comprehensive characterisation of the wheat TPE for north-eastern Australia (Chenu et al., 2011; Bustos-Korts et al., 2021) and so we provided some additional interpretation of GxE interactions for yield related to water availability and the incidence of drought and their influences on the genetic correlation *r*_*g*(*MET,TPE*)_ in terms of the more recent TPE characterisation (Figures 1 and 2).

### Future research

The extended form of the breeder’s equation is particularly relevant as a framework for the design of breeding strategies to target climate resiliency to address the impacts of climate change on the environmental composition of the short, medium and long-term future diverse geographical TPEs expected for our global agricultural systems (Chapman et al. 2012; IPCC, 2021; Cooper and Messina, 2022). Future work will explore developments and other applications of the extended breeder’s equation to assist design of prediction-based breeding programs to tackle the effects of climate change, where it is expected that frequencies of environment-types within the TPE will change with time (Chapman et al., 2012; Lobell et al., 2015; Hammer et al. 2020; Snowdon et al., 2020; Cooper et al., 2021; IPCC, 2021; Bustos-Korts et al., 2021; Cooper and Messina, 2022).

## Author Contributions

MC conceived and wrote the manuscript. Ideas that contributed to the manuscript came from collaborative research conducted by MC, CG, CM, TT, OP. All authors read and provided comments that contributed to the manuscript.

## References

Araus, J.L., and Cairns, J.E. (2014). Field high-throughput phenotyping, the new frontier in crop breeding. Trends in Pl. Sci. 19, 52–61.

Basford, K.E., and Cooper, M. (1998). Genotype x environment interactions and some considerations of their implications for wheat breeding in Australia. Aust. J. Agric. Res. 49, 153–174.

Blum, A. (1988). Plant Breeding for Stress Environments. Boca Raton, FL, USA: CRC Press.

Braun, H.-J., Atlin, G., and Payne, T. (2010). “Multi-location testing as a tool to identify plant response to global climate change,” in Climate change and crop production, ed. M.P. Reynolds (Wallingford, UK: CAB International), 115–138.

Brennan, P.S., Byth, D.E., Drake, D.W., De Lacy, I.H., and Butler, D.G. (1981). Determination of the location and number of test environments for a wheat cultivar evaluation program. Aust. J. Agric. Res. 32, 189–201.

Bustos-Korts, D., Boer, M.P., Chenu, K., Zheng, B., Chapman, S., van Eeuwijk, F. (2021). Genotype specific P-spline response surfaces assist interpretation of regional wheat adaptation to climate change. In silico Plants 3. doi: 10.1093/insilicoplants/diab018

Ceccarelli, S. (1989). Wide adaptation: How wide? Euphytica 40, 197–205.

Ceccarelli, S. (1994). Specific adaptation and breeding for marginal conditions. Euphytica 77, 205–219.

Ceccarelli, S., Grando, S. (2020). Evolutionary plant breeding as a response to the complexity of climate change. iScience 23. doi: 10.1016/j.isci.2020.101815

Chapman, S.C., Chakraborty, S., Dreccer, M.F., and Howden, S.M. (2012). Plant adaptation to climate change – opportunities and priorities in breeding. Crop Pasture Sci. 63, 251–268.

Chenu, K., Cooper, M., Hammer, G.L., Mathews, K.L., Dreccer, M.F., and Chapman, S.C. (2011). Environment characterization as an aid to wheat improvement: interpreting genotype-environment interactions by modelling water-deficit patterns in North-Eastern Australia. J. Exp. Bot. 62, 1743–1755.

Cobb, J.N., Juma, R.U., Biswas, P.S., Arbelaez, J.D., Rutkoski, J., Atlin, G., et al. (2019). Enhancing the rate of genetic gain in public-sector plant breeding programs: lessons from the breeder’s equation. Theor. Appl. Genet. 132, 627–645.

Comstock, R.E. (1996). Quantitative genetics with special reference to plant and animal breeding. Ames, IA: Iowa State University Press.

Comstock, R.E., and Moll, R.H. (1963). “Genotype-Environment interactions” in Statistical genetics and plant breeding, eds W.D. Hanson and H.F. Robinson, (Washington, D.C., USA: Publication 982, National Academy of Sciences – National Research Council), 164–196.

Cooper, M., and DeLacy, I.H. (1994). Relationships among analytical methods used to study genotypic variation and genotype-by-environment interaction in plant breeding multi-environment experiments. Theor. Appl. Genet. 88, 561–572.

Cooper, M., Gho, C., Leafgren, R., Tang T., and Messina, C. (2014b). Breeding drought-tolerant maize hybrids for the US corn-belt: discovery to product. J. Exp. Bot. 65, 6191–6204.

Cooper, M., and Messina, C.D. (2022). Breeding crops for drought-affected environments and improved climate resilience. Plant Cell 2022, doi: 10.1093/plcell/koac321

Cooper, M., Messina, C.D., Podlich, D., Totir, L.R., Baumgarten, A., Hausmann, N.J., et al. (2014a). Predicting the future of plant breeding. Complementing empirical evaluation with genetic prediction. Crop Pasture Sci. 65(4), 311–336.

Cooper, M., Stucker, R.E., DeLacy, I.H., and Harch, B.D. (1997). Wheat breeding nurseries, target environments, and indirect selection for grain yield. Crop Sci 37, 1168–1176.

Cooper, M., Tang, T., Gho, C., Hart, T., Hammer, G., and Messina, C. (2020). Integrating genetic gain and gap analysis to predict improvements in crop productivity. Crop Sci 60, 582–604.

Cooper, M., Voss-Fels, K.P., Messina, C.D., Tang, T., and Hammer, G.L. (2021). Tackling GxExM interactions to close on-farm yield-gaps: creating novel pathways for crop improvement by predicting contributions of genetics and management to crop productivity. Theor. Appl. Genet. 134, 1625–1644.

Cooper, M., Woodruff, D.R., Eisemann, R.L., Brennan, P.S., and DeLacy, I.H, (1995). A selection strategy to accommodate genotype-by-environment interaction for grain yield of wheat: managed-environments for selection among genotypes. Theor. Appl. Genet. 90, 492–502.

Cooper, M., Woodruff, D.R., Phillips, I.G., Basford, K.E., and Gilmour, A.R, (2001). Genotype-by-management interactions for grain yield and grain protein concentration of wheat. Field Crops Res. 69, 47–67.

de los Campos, G., Pérez-Rodriguez, P, Bogard, M., Gouache, D., Crossa, J. (2020). A data-driven simulation platform to predict cultivars’ performances under uncertain weather conditions. Nature Comm. 11, 4876. doi: 10.1038/s41467-020-18480-y

Diepenbrock, C., Tang, T., Jines, M., Technow, F., Lira, S., Podlich, D., et al. (2021). Can we harness digital technologies and physiology to hasten genetic gain in U.S. maize breeding? Plant Physiology 188(2), 1141–1157.

Duvick, D.N. (2001). Biotechnology in the 1930s: the development of hybrid maize. Nature Reviews, Genetics 2, 69–74.

Duvick, D.N., Smith, J.S.C., and Cooper, M. (2004). Long-term selection in a commercial hybrid maize breeding program. Plant Breeding Reviews 24, 109–151.

Falconer, D.S. (1952). The problem of environment and selection. American Naturalist 86, 293–298.

Gaffney, J., Schussler, J., Löffler, C., Cai, W., Paszkiewicz, S., Messina, C., et al. (2015). Industry-scale evaluation of maize hybrids selected for increased yield in drought-stress conditions of the US corn belt. Crop Sci. 55, 1608–1618.

Goldman, I.L. (2000). Prediction in plant breeding. Plant Breeding Reviews 19, 15–40.

González-Barrios, P., Díaz-García L., and Gutiérrez, L. (2019). Mega-Environmental Design: Using Genotype x Environment Interaction to Optimize Resources for Cultivar Testing. Crop Sci. 59, 1899–1915.

Hajjarpoor, A., Kholová, J., Pasupuleti, J., Soltani, A., Burridge, J., Degala, S.B., et al. (2021). Environmental characterization and yield gap analysis to tackle genotype-by-environment-by-management interactions and map region-specific agronomic and breeding targets in groundnut. Field Crops Res. 267, 108160. doi: 10.1016/j.fcr.2021.108160

Haldane, J.B.S. (1947). The interaction of nature and nurture. Ann. Eugenics. 13, 197–205.

Hallauer, A.R., and Miranda, J.B.F. (1988). Quantitative Genetics in Maize Breeding, Second Edition. Ames, IA, USA: Iowa State University Press.

Hammer, G.L., McLean, G., van Oosterom, E., Chapman, S., Zheng, B., Wu, A., et al. (2020). Designing crops for adaptation to the drought and high-temperature risks anticipated in future climates. Crop Sci 60, 605–621.

Heffner, E.L., Sorrells, M.E., and Jannink, J.L. (2009). Genomic selection for crop improvement. Crop Sci. 49, 1–12.

IPCC, (2021). Climate Change 2021: The Physical science basis. Contribution of Working Group I to the sixth assessment report of the Intergovernmental Panel on Climate Change, Cambridge University Press, UK.

Jarquín, D., Crossa, J., Lacaze, X., Du Cheyron, P., Daucourt, J., Lorgeou, J., et al. (2014). A reaction norm model for genomic selection using high-dimensional genomic and environmental data. Theor. Appl. Genet. 127, 595–607.

Kholová, J., McLean, G., Vadez, V., Craufurd P., and Hammer, G.L. (2013). Drought stress characterization of post-rainy season (rabi) sorghum in India. Field Crops Res. 141, 38–46.

Kholová, J., Urban, O., Cock, J., Arcos, J., Arnaud, E., Aytekin, D., et al. (2021). In pursuit of a better world: crop improvement and the CGIAR. J. Exp. Bot. 72(14), 5158–5179.

Langstroff, A., Heuermann, M.C., Stahl, A., and Junker, A. (2022). Opportunities and limits of controlled-environment plant phenotyping for climate response traits. Theor. Appl. Genet. 135, 1–16.

Lobell, D.B., Hammer, G.L., Chenu, K., Zeng, B., McLean G., and Chapman, S.C. (2015). The shifting influence of drought and heat stress for crops in northeast Australia. Global Change Biol. 21, 4115–4127.

Lush, J.L. (1937). Animal Breeding Plans. Ames, IA: Iowa State University Press.

Messina, C.D., Ciampitti, I., Berning, D., Bubeck, D., Hammer, G.L., and Cooper M. (2022a). Sustained improvement in yield stability accompanies maize yield increase in temperate environments. Crop Sci. 62, 2138–2150.

Messina, C.D., Tang, T., Truong, S.K., McCormick, R.F., Technow, F., Powell, O., et al. (2022b). Crop improvement for circular agricultural systems. J. ASABE 65, 491–504.

Messina, C.D., Technow, F., Tang, T., Totir, R., Gho C., and Cooper, M. (2018). Leveraging biological insight and environmental variation to improve phenotypic prediction: Integrating crop growth models (CGM) with whole genome prediction (WGP). Eur. J. Agron. 100, 151–162.

Meuwissen, T.H., Hayes, B.J., and Goddard, M.E. (2001). Prediction of total genetic value using genome-wide dense marker maps. Genetics 157(4), 1819–1829.

Millet, E.J., Kruijer, W., Couple-Ledru, A., Prado, S.A., Cabrera-Bosquet, L., Lacube, S., et al. (2019). Genomic prediction of maize yield across European environmental conditions. Nat. Genet. 51, 952–956.

Nyquist, W.E., and Baker, R.J. (1991). Estimation of heritability and prediction of selection response in plant populations. Crit. Rev. Pl. Sci. 10(3), 235–322.

Resende, R.T., Piepho, H.P., Rosa, G.J.M., Silva-Junior, O.B., e Silva, F.F., de Resende, M.D.V., et al. (2021). Enviromics in breeding: applications and perspectives on envirotypic-assisted selection. Theor. Appl. Genet. 134, 95–112.

Rogers, A.R., Dunne, J.C., Romay, C., Bohn, M., Buckler, E.S., Ciampitti, I.A., et al. (2021). The importance of dominance and genotype-by-environment interactions on grain yield variation in a large-scale public cooperative maize experiment. G3 Genes Genomes and Genetics 11(2), doi: 10.1093/g3journal/jkaa050

Ronanki S., Pavlík J., Masner J., Jarolímek J., Stočes M., Subhash D., et al. (2022). An APSIM-powered framework for post-rainy sorghum-system design in India. Field Crops Res. 277, 108422, doi: 10.1016/j.fcr.2021.108422

Smith, A.B., Cullis B.R., and Thompson, R. (2005). The analysis of crop cultivar breeding and evaluation trials: an overview of current mixed model approaches. J. Agr. Sci. 143, 449–462.

Smith, A., Ganesalingam, A., Lisle, C., Kadkol, G., Hobson K., and Cullis, B. (2021a). Use of contemporary groups in the construction of multi-environment trial datasets for selection in plant breeding programs. Front. Plant Sci. 11, 623586, doi: 10.3389/fpls.2020.623586

Smith, A., Norman, A., Kuchel, H., and Cullis, B. (2021b). Plant variety selection using interaction classes derived from factor analytic linear mixed models: Models with independent variety effects. Front. Plant Sci. 12, 737462, doi: 10.3389/fpls.2021.737462

Snowdon, R.J., Wittkop, B., Chen, T.-W., Stahl, A. (2020). Crop adaptation to climate change as a consequence of long-term breeding. Theor. Appl. Genet. 134, 1613–1623.

van Eeuwijk, F., Bustos-Korts, D., Millet, E.J., Boer, M., Kruijer, W., Thompson, A. et al. (2019). Modelling strategies for assessing and increasing the effectiveness of new phenotyping techniques in plant breeding. Plant Sci. 282, 23–39.

van Eeuwijk, F.A., Cooper, M., DeLacy, I.H., Ceccarelli, S., and Grando, S. (2001). Some vocabulary and grammar for the analysis of multi-environment trials, as applied to the analysis of FPB and PPB trials. Euphytica 122, 477–490.

Varshney, R.K., Bohra, A., Yu, J., Graner, A., Zhang, Q., and Sorrells, M.E. (2021). Designing future crops: Genomics-assisted breeding comes of age. Trends Plant Sci 26, 631–649.

Voss-Fels, K.P., Cooper M., and Hayes, B.J. (2019). Accelerating crop genetic gains with genomic selection. Theor. Appl. Genet. 132, 669–686.

Xiong, W., Reynolds, M.P., Crossa, J., Schulthess, U., Sonder, K., Montes, C., et al. (2021). Increased ranking change in wheat breeding under climate change. Nat. Plants 7, 1207–1212.

